# Efficient parameter estimation for ODE models of cellular processes using semi-quantitative data

**DOI:** 10.1101/2024.01.26.577371

**Authors:** Domagoj Dorešić, Stephan Grein, Jan Hasenauer

## Abstract

Quantitative dynamical models facilitate the understanding of biological processes and the prediction of their dynamics. The parameters of these models are commonly estimated from experimental data. Yet, experimental data generated from different techniques do not provide direct information about the state of the system but a non-linear (monotonic) transformation of it. For such semi-quantitative data, when this transformation is unknown, it is not apparent how the model simulations and the experimental data can be compared. Here, we propose a versatile spline-based approach for the integration of a broad spectrum of semi-quantitative data into parameter estimation. We derive analytical formulas for the gradients of the hierarchical objective function and show that this substantially increases the estimation efficiency. Subsequently, we demonstrate that the method allows for the reliable discovery of unknown measurement transformations. Furthermore, we show that this approach can significantly improve the parameter inference based on semi-quantitative data in comparison to available methods. Modelers can easily apply our method by using our implementation in the open-source Python Parameter EStimation TOolbox (pyPESTO).

## Introduction

The use of mechanistic mathematical models has greatly contributed to the understanding of biological processes at the cellular [13, 25], patient [5, 11] and population level [9, 31]. In particular, mechanistic ordinary differential equation (ODE) models are used for a broad spectrum of applications, ranging from cellular signaling, metabolism, and gene regulation over pharmacokinetics and -dynamics to the spread of diseases. However, ODE models often contain parameters that cannot be measured directly. Instead, the parameters have to be estimated from experimental data [17]. This is commonly achieved by numerical optimization of an objective function, which quantifies how well the model simulations fit the given experimental data, such as the likelihood function.

The experimental data used for parameter estimation are collected using a broad spectrum of experimental techniques. For example, early studies in the field of systems biology employed well-calibrated Western blot experiments and performed an in-depth assessment of the mapping of concentration to measured intensities [15]. In this case, the data were ensured to fall within the linear regime of the experimental technique, and often even absolute quantification was performed. However, many, even state–of–the–art, measurement techniques do not ensure a linear relationship between the abundance of the biochemical quantities of interest and the measured output (Fig. 1). Well-known examples include fluorescence microscopy data such as Förster resonance energy transfer (FRET) data [1], optical density (OD) measurement [29] and imaging data for certain stainings [18]. In addition, many experimental techniques suffer from lower limits of detection and/or saturation effects.

**Figure 1:**
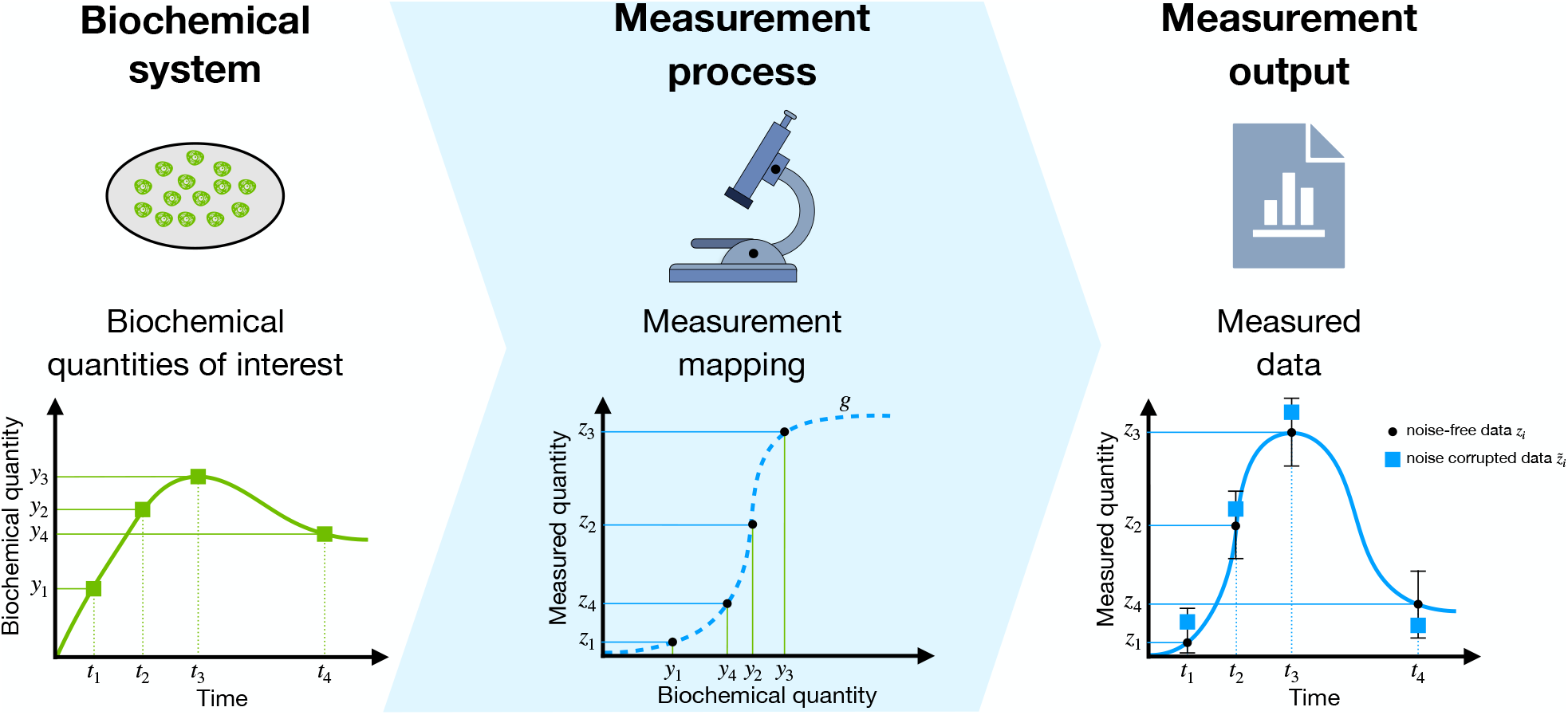
A non-linear measurement mapping. (A) True values of a biochemical quantity of interest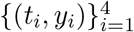. (B) A measurement process can introduce unknown non-linear data mappings (dashed blue line). In that case, a mapping function transforms the biochemical quantities 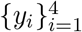 and yields non-linearly mapped measured quantities 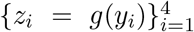. (C) The measurement quantities 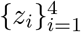 are corrupted by noise, resulting in a noise-corrupted dataset 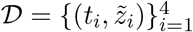.

Quantitative data are easy to use for the parameterization of ODE models and the same holds for data that are collected in the linear regime of measurement devices. This is showcased in a large number of published articles (see, e.g. [10] for a collection of models and datasets). In fact, there are custom methods for experimental data for which a linear mapping with unknown scaling and offset parameters can be assumed [16, 22]. If the linearity assumption is not fulfilled, it is usually assumed that the mapping from biochemical quantities of interest to measured output is monotone. This monotonicity ensures that the ordering is preserved and allows the use of approaches for ordinal data, such as the optimal scaling method [27]. For ODE models, this approach has recently been accelerated by using a reformulation of the optimization problem [23] and gradient information [24]. However, in this approach, all quantitative information is discarded and the defined objective function is not based on probabilistic grounds, disallowing any uncertainty analysis.

In this manuscript, we introduce a spline-based approach to use semi-quantitative data – which are obtained using an experimental technique with a non-linear but monotone mapping – for parameter estimation. We assume that the measurement mappings are monotone, as is frequently observed in experiments. The method reconstructs the unknown mapping function using a statistically coherent formulation that facilitates uncertainty analysis. We demonstrate the credibility of the proposed approach as a tool for uncovering measurement mapping shapes. To illustrate the parameter inference capabilities of the method, we benchmark its performance with a collection of published models. Furthermore, we derive formulas for the analytical calculation of the gradients of the objective function in hierarchical optimization. To evaluate this optimization framework, we compare its efficiency with alternative approaches.

## Methods

### Mechanistic modeling of biological systems

We consider models of biological processes based on systems of ODEs:

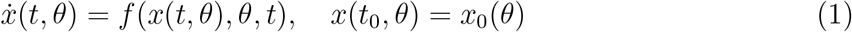

in which the temporal evolution of the state variables 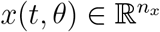 is determined by the vector field 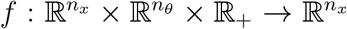 with the unknown mechanistic model parameters 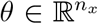. The parameters usually consist of kinetic rate constants and initial species conditions. The measured properties of a model are its observables 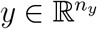,

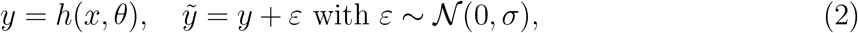

in which 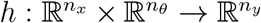 denotes the observation map which models the dependence of the observables on the model state variables and unknown mechanistic parameters, 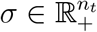 is a noise parameter, and 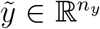 are noise-corrupted measurements. The dimensionalities of the state, parameter, and observable vector are denoted by *n*_*x*_, *n*_*θ*_, and *n*_*y*_, respectively. The number of time points is denoted by *n*_*t*_.

### Linear semi-quantitative (relative) observables

Most measurement techniques provide only relative information on the biochemical quantity of interest. In this case, to obtain values comparable to the measured quantities, those observables need to be rescaled by scaling factors *a* and offsets *b*. Most often, an additive Gaussian distributed noise model is assumed. Then the full relationship between measured and biochemical quantities of a relative observable is given by:

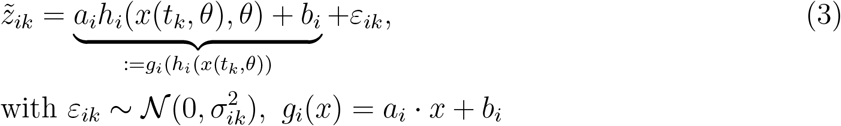

in which *i* is the observable index, *k* is the time index, and *g*_*i*_ : ℝ → ℝ is an affine measurement mapping from the *i*-th observable, *h*_*i*_(*x*(*t*_*k*_, *θ*)), to its measurement. The scaling factors, offsets, and noise parameters of the *i*-th observable are denoted as *a*_*i*_ ∈ ℝ, *b*_*i*_ ∈ ℝ, and 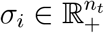, respectively. These parameters are often unknown and need to be estimated along with all other unknown parameters of the model.

### Non-linear semi-quantitative observables

In some cases, the measurement process induces a non-linear mapping between the biochemical quantities of interest and the measured quantities. Assuming an additive Gaussian distributed noise model the relationship is given by:

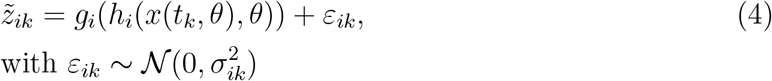

in which *g*_*i*_ : ℝ → ℝ is a non-linear measurement mapping from the *i*-th observable, *h*_*i*_(*x*(*t*_*k*_, *θ*)), to its measurement. The form and parameterization of non-linear measurement mappings *g*_*i*_(*h*_*i*_(*x, θ*)) are application-dependent. We provide examples in the result section. Measurement mappings are often unknown and need to be modeled in some way.

As this study considers the data-driven uncovering of measurement mappings, we select a class of approximations. Specifically, we consider the approximation of measurement mappings *g*_*i*_(*h*_*i*_(*x, θ*)) with monotone, piece-wise linear splines 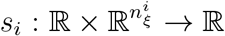, in which *I* is the observable index. For the simplicity of further calculation, we parameterize the splines using the differences between the heights of neighboring spline knots 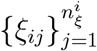:

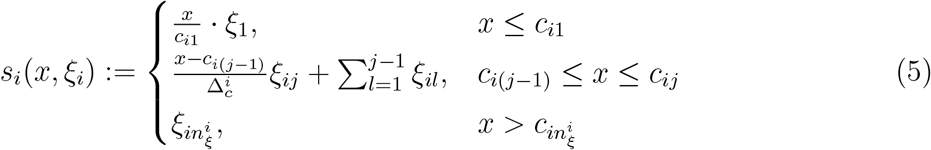

in which 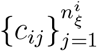 are the knot bases, and 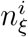 is the number of spline knots for the *i*-th observable. Since *g*_*i*_ is monotone, we constrain the spline parameters to be positive *ξ*_*i,j*_ *≥* 0 for all 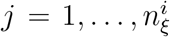. We regularized the spline by adding a penalty term to the objective function to promote linearity (Fig. 2B), which greatly improved the convergence of the estimation. For details on the definition of the spline, the distribution of the knot bases, and spline regularization, we refer to the first section of the supplementary materials.

**Figure 2:**
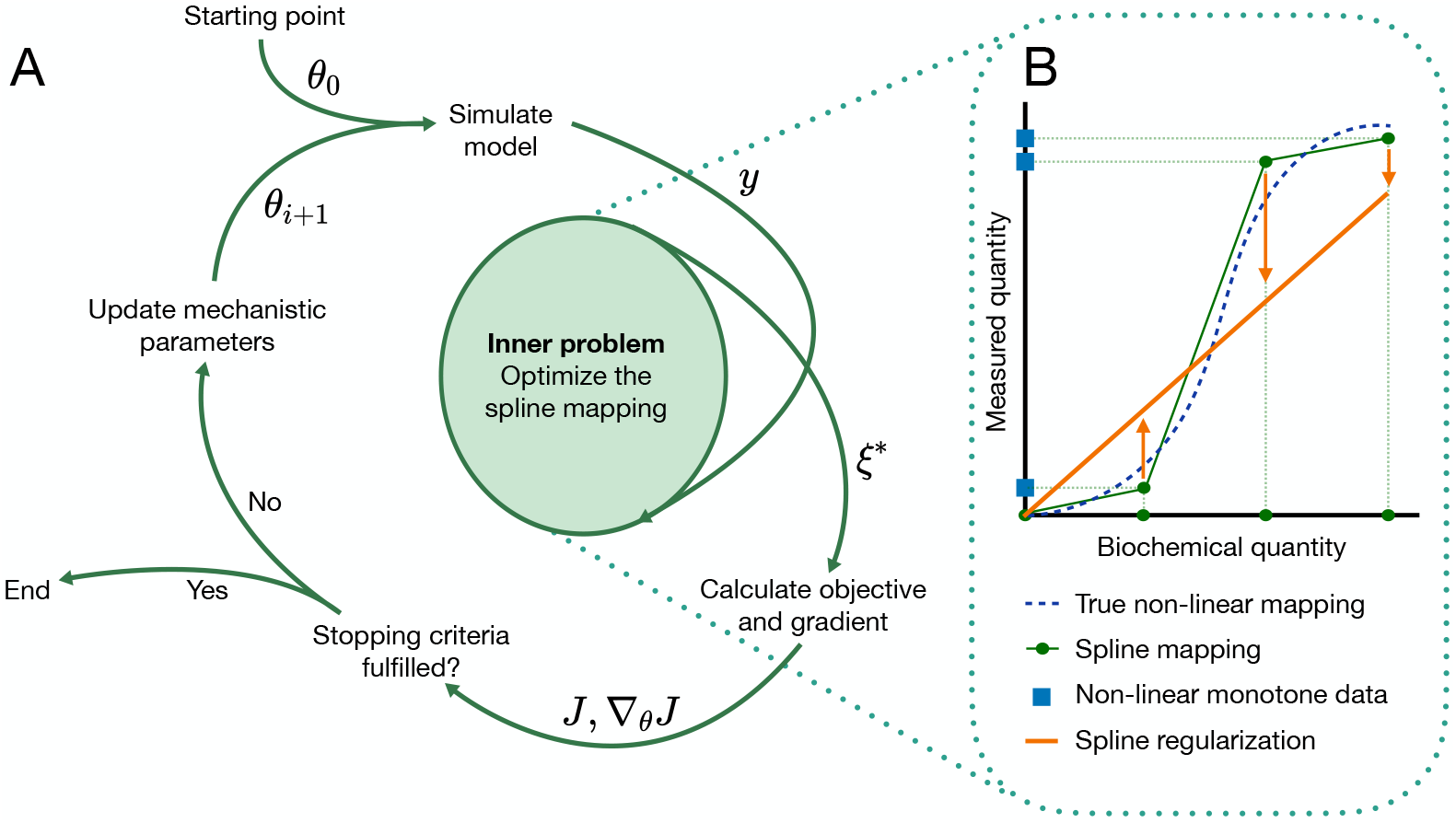
Illustration of the spline estimation approach. (A) Model mechanistic parameters *θ* are iteratively updated during parameter estimation. For each vector of trial parameters, the model is simulated to obtain the simulation *y*. Then, the spline parameters *ξ*\* are optimized and used to calculate the objective function *J* and its gradient *∇*_*θ*_*J*. These are then passed on to obtain the next trial parameter vector, or the optimization is halted. (B) The spline (green) enables mapping of the simulation of the model *y* (biochemical quantities) to the measurement axis. This allows for the definition of a likelihood objective function. In the inner problem, this objective function is minimized with respect to the spline parameters to obtain optimal spline parameters *ξ*\*. The spline is additionally regularized by the distance to the linear mapping (orange).

The spline allows us to link the measured and biochemical quantities of the non-linear monotone observable:

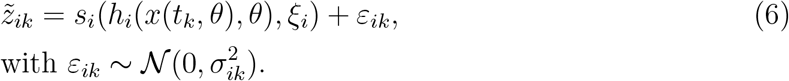

The model dataset 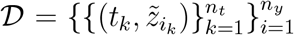 consists of observations of all model observables at time-points 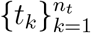. We denote the dataset of the *i*-th observable as 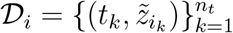.

### Parameter estimation

For a dataset 𝒟 consisting of independent observations of quantitative, linear semi-quantitative, and/or non-linear semi-quantitative observables, we can define the negative log-likelihood objective function as:

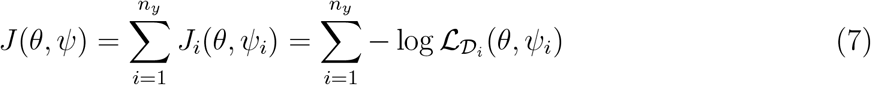

in which *ψ*_*i*_ are the observable parameters of the *i*-th observable: for a relative observable these are scaling *a*_*i*_ and offset *b*_*i*_, while for a non-linear semi-quantitative observable they are the spline parameters *ξ*_*i*_. Minimizing the objective function, we obtain the maximum likelihood estimate (MLE): 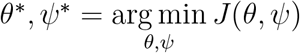.

### Hierarchical optimization problem and analytical gradients

The objective function minimization can be executed jointly in all observable models and parameters. However, this leads to a high-dimensional optimization problem and long computation times. Alternatively, the optimization problem can be separated hierarchically (Fig. 2A):

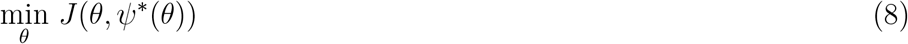

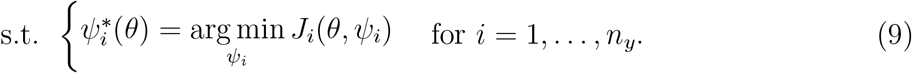

In the outer optimization problem (8), we estimate the mechanistic parameters, and in the inner optimization problems (9) we estimate the observable parameters *ψ*_*i*_ of each observable. For non-linear semi-quantitative observables, the inner problem is additionally constrained by the positivity of spline parameters. Since the objective function can be additively separated into components 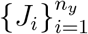 depending on the observable parameters of a single observable *ψ*_*i*_, the inner optimization problem is a set of *n*_*y*_ small inner optimization problems (9). For relative observables, these inner problems can be solved analytically [22]. For non-linear semi-quantitative observables, they still need to be numerically minimized, but the inner problems are convex (Theorem 1 in the supplementary materials) and thus easy to minimize. Furthermore, the gradient of the objective function can be calculated analytically. To show this, let the *i*-th observable be non-linear semi-quantitative for some *i* ∈ {1, …, *n*_*y*_}. As the spline parameters are estimated hierarchically, their optimal values 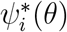 depend on the trial mechanistic parameter vector *θ*. This adds another term to the analytical gradient calculation of the outer optimization

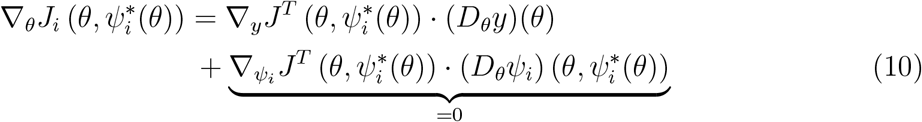

in which (*D*_*x*_*f*)(*x*\*) is the Jacobian of a vector-valued function *f* with respect to variable vector *x* evaluated at a point *x*\*. The first term of the gradient is standard and can be analytically calculated using forward sensitivity analysis. The second term is the result of the dependency mentioned. However, in the supplementary materials, we prove that this term is always 0 (Theorem 2). Thus, the gradient can be analytically computed.

### Confidence region of a parameter vector *θ*

Here, we define what it means for a parameter vector to lie in a confidence region of a certain significance. We do this using the likelihood-ratio test in which we define the corresponding test statistic as

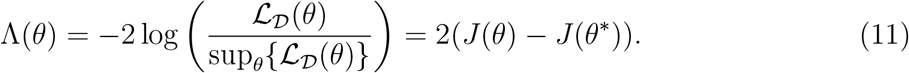

In the asymptotic case of a large number of data points, the test statistic converges to a chi-square distribution 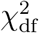 with df = *n*_*θ*_ degrees of freedom (see [30] for more details). Then we define the confidence region of significance *α* as

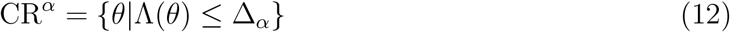

in which Δ_*α*_ is the *α*-th percentile of the 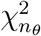 distribution.

### Benchmark models

For the evaluation of the proposed method, we consider one exemplary model and four published models that were previously developed and calibrated for different biological systems (Table 1). As the published models originally did not contain non-linear semi-quantitative observables, we generated synthetic data at the same time points, chose non-linear monotone measurement mappings, applied them to the observables, and corrupted them with the same type of noise as in the original model. For details on the synthetic data generation, chosen non-linear measurement mappings, and model structure, we refer to the second section of the supplementary material.

**Table 1:** Benchmark models.

| Model | $n_x$ | $n_\theta$ | $n_y$ | $\vert \mathcal{D}\vert $ | Description |
| --- | --- | --- | --- | --- | --- |
| T1 | 2 | 4 | 1 | 12 | FRET probe activation [1] |
| M1 | 8 | 6 | 3 | 48 | STAT5 dimerization [2] |
| M2 | 7 | 9 | 1 | 23 | Infectious disease dynamics [19] |
| M3 | 8 | 18 | 1 | 58 | Transcriptional regulation [4] |
| M4 | 14 | 18 | 8 | 205 | IL13-induced signaling [20] |

Note: By $|\mathcal{D}|$ we denote the cardinality of the dataset.

## Results

### The proposed method uncovers measurement mapping for FRET probe activation

To illustrate an application of the proposed method, we have applied it to a FRET probe activation model introduced by [1]. In general, the transition of inactive FRET probes *P* to an active state *P*\* can be represented by the scheme in Fig. 3A. The quantity of interest in this model is the ratio of activated probes to total probes *P*\**/P*_*TOT*_. The most common way to measure this value is through a measurement technique called ratiometric imaging. Cells are exposed to excitation light from the donor channel, and then fluorescence emission is divided into donor and acceptor channels. The output of ratiometric imaging, R, is the intensity in the acceptor channel, *I*_*A*_, divided by the intensity in the donor channel, *I*_*D*_. Previous studies have shown that this measured R value can have a highly non-linear relationship to the fraction of active FRET probes [1] (Fig. 3A).

**Figure 3:**
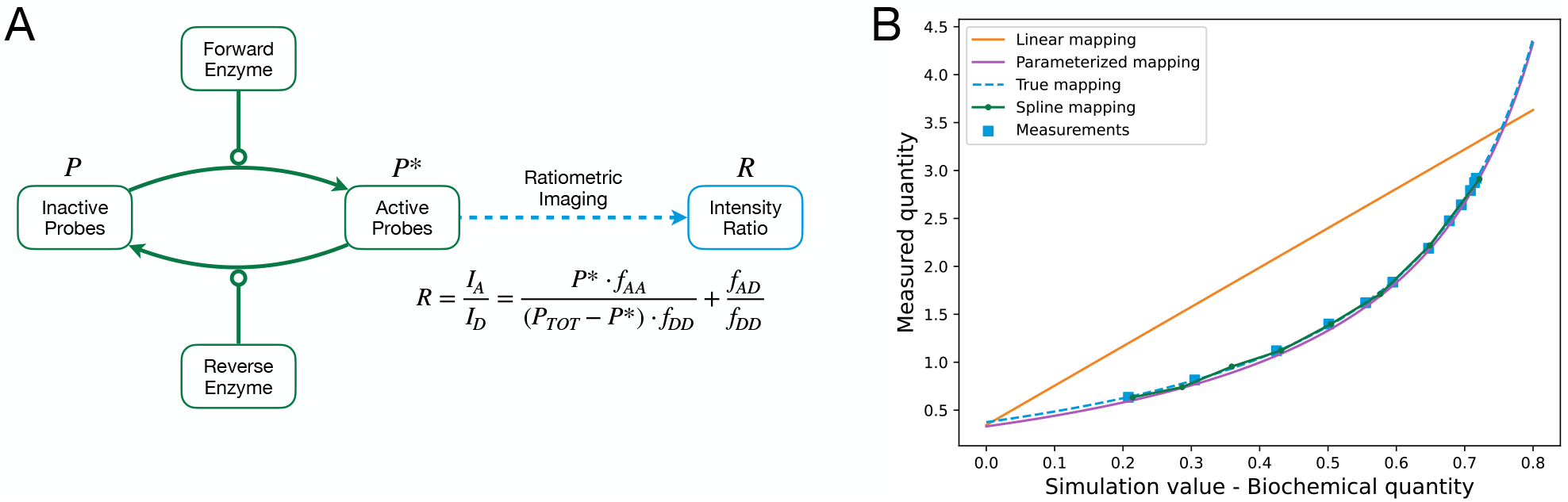
A model of FRET probe activation. (A) A forward enzyme catalyzes the activation of inactive FRET probes *P*, and a reverse enzyme catalyzes the conversion of an active probe *P*\* into the inactive state. Active probe concentration can be observed via ratiometric imaging. The measurement mapping of this process is highly non-linear. *f*_*AA*_ and *f*_*AD*_ are fractions of the acceptor and donor emissions that the acceptor channel captures, respectively. *f*_*DD*_ is the fraction of donor emissions that the donor channel captures. (B) Comparison of the estimation of the measurement mapping using a linear function, proposed spline mapping, and parameterization of the true mapping. For all three models, we performed 1000 local optimizations. Depicted are the estimated mappings closest to the true mapping of starts with mechanistic parameters in the 95% confidence region. We show the synthetic noise-corrupted data used in all model optimizations in blue squares.

One approach of modeling this non-linear mapping is to parameterize a function of a similar shape and to estimate its parameters. For FRET probe activation it has been shown that the relation between state variables and measurement is of the form:

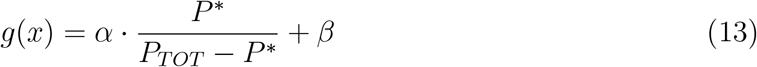

with experiment- and probe-specific parameters *α* and *β*. However, this requires prior knowledge of the shape of the measurement mapping. Without such prior knowledge, the measurement mapping has to be inferred. A simple and easy-to-implement approach is to assume that the mapping is linear. This linear approximation can be sufficiently correct if the measurement region is locally linear. For highly non-linear measurement mappings, this is not true, so one has to resort to more flexible approaches such as spline estimation.

To evaluate how well the three modeling approaches can recover the true measurement mapping, we performed 1000 local optimizations for each and chose the best measurement mapping estimates in the 95% confidence region (Fig. 3 B). We found that the reconstruction using spline estimation agrees well with the true measurement mapping. Indeed, it yields similar results to using parametric representations with unknown parameters. In contrast, a linear model for the measurement mapping proved to be insufficiently flexible and resulted in biased reconstruction of the measurement mapping.

Overall, our assessment revealed that, unlike a simple linear approximation for the measurement mapping, a spline-based approximation enables the reconstruction of non-linear mappings as observed for FRET probe activation.

### The spline estimation approach as a tool for uncovering measurement mapping shapes

In the previous subsection, we have shown that the estimation of an unknown measurement mapping using a spline can yield results similar to the estimation of a parametric representation. Yet, we only considered a point estimate and did not assess the reliability of the reconstruction. To determine whether the proposed approach provided statistically coherent estimates, we considered model M1 with measurement mappings of various shapes across observables (Fig. 4A-C). Using the resulting dataset, we performed a multi-start optimization (10^3^ runs) to obtain optimal parameters and Markov chain Monte Carlo sampling using an adaptive Metropolis-Hastings algorithm (10^5^ iterations). The resulting chain was thinned by a factor of 500. We computed the optimal spline for each of the remaining samples and, with them, constructed the credibility intervals of the optimal spline mappings.

**Figure 4:**
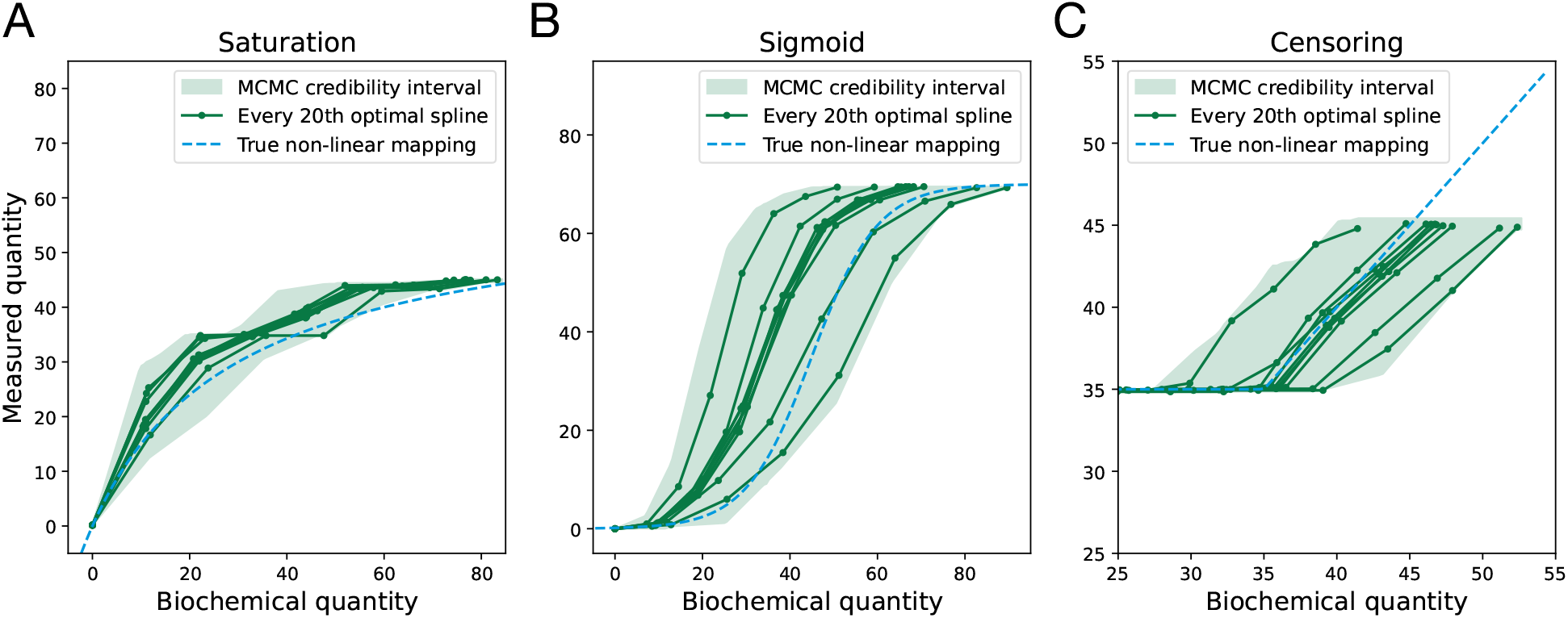
Credibility intervals of the estimated spline mappings. MCMC sampling of the model M1. The model contains different synthetic non-linear measurement mappings for each of its observables (dashed blue). Splines were estimated for each 500th sample of the MCMC run with which we constructed the credibility intervals of the estimated spline mappings (light green). For visibility, we show only each 20th estimated spline (dark green).

The inspection of the results confirmed that the optimal splines are qualitatively similar to the measurement mappings used for data generation. Furthermore, more importantly, the measurement mappings used for data generation lie within the credibility intervals. This showcases the reliability of the method as a tool for discovering curve shapes of unknown measurement mappings.

### Hierarchical optimization and analytical gradients increase the estimation efficiency

As the method provides reliable estimates for the mappings, we turn to the assessment of the computational cost, which is of high practical relevance. Here, we also wanted to evaluate the impact of (i) reformulation as a hierarchical problem and (ii) the availability of analytical gradients. For this assessment, we considered the published models M1 to M4 with synthetic data with a range of different measurement mappings, as detailed in the second section of the supplementary material. For all models, we performed 1000 local optimizations with equal start points across approaches. We then determined the overall computational cost, the number of function evaluations, and the number of converged starts per computation time.

We found that in general, the proposed hierarchical approach with analytical gradients achieves the best performance (Fig. 5). This appears to be mostly related to a reduction in the computation time, respectively, the number of function evaluations (Fig. 5A, B), while the number of converged starts remains rather similar (Fig. 5C). The number of converged starts per computation time is, for the hierarchical approach with analytical gradients, at least twice as high for the other approaches (Fig. 5D). Interestingly, a hierarchical approach without gradient information does not perform well, and is worse than the non-hierarchical approach with gradient information for all models.

**Figure 5:**
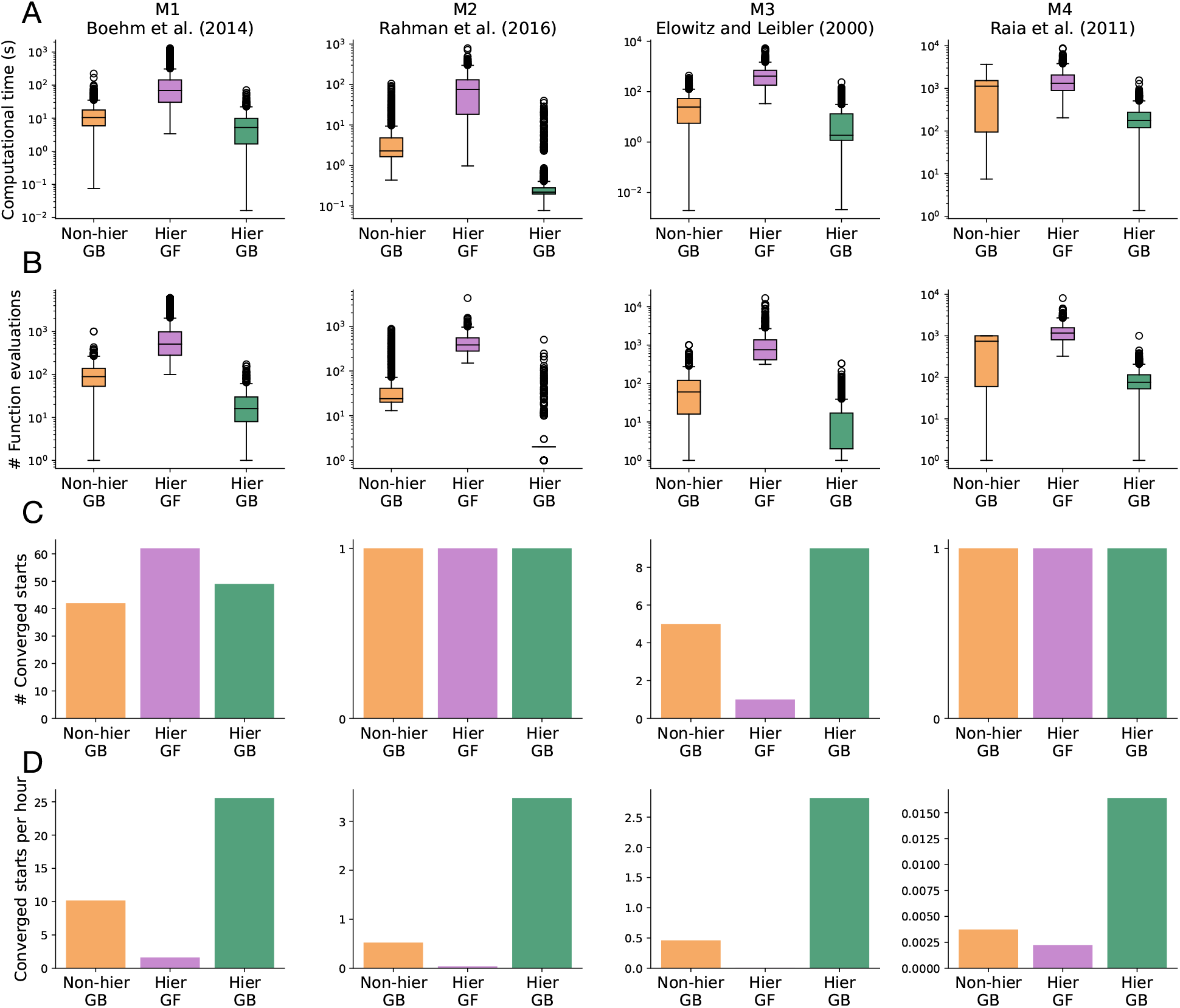
Evaluation of the gradient-based non-hierarchical, gradient-free hierarchical, and gradient-based hierarchical estimation approaches. Models M1 - M4 are shown from left to right. (A) Comparisons of computation time. (B) Comparisons of the number of function evaluations. (C) Comparisons of the number of converged starts. Converged starts are defined as the starts with estimated mechanistic parameters within the 95% confidence region. (D) Comparison of the number of converged starts per CPU hour.

For the proposed hierarchical approach with analytical gradients, the number of converged starts per CPU hour was on average *≈* 7.96. As the spread between models was large, this finding clearly suggests that the approach is computationally tractable.

### Spline approach improves the parameter inference of models with unknown measurement mappings

Our proposed method provides reliable estimates of measurement mappings. Here, we examine whether this leads to good estimates of the mechanistic model parameters. Apart from spline estimation, a generally applicable approach to the integration of semi-quantitative data into parameter estimation is linear estimation of measurement mappings. Thus, we compare the parameter inference of these two approaches in a realistic setting. In addition, for reference, we include the approach of discarding data with unknown measurement mappings. We performed 1000 local optimizations for the application examples M1-M4. We evaluated the impact of an increasing number of unknown measurement mappings by turning quantitative observables into semi-quantitative observables. As a parameter inference metric, we use the mean L2 distance of the estimated to the true mechanistic parameters normalized by the number of mechanistic parameters. For details of the study setting, we refer to the fourth section of the supplementary materials.

The spline estimation outperforms other approaches (Fig. 6). In general, linear estimation has a stronger bias than variance. We observe this primarily for model M4, as the linear estimation has the smallest standard deviation between approaches (Fig. 6 D). In some cases, this even causes the linear estimation to perform worse than the approach of discarding data with unknown mappings. In contrast, the higher flexibility of the spline estimation allows for the general attainment of better parameter estimates. This is the case even for model M4 with eight unknown measurement mappings, for which the spline estimation adds seven times more parameters than the linear estimation. For a small number of unknown measurement mappings, the spline estimation can perform almost equally well as the model with completely known measurement mappings. This showcases that the proposed method yields good estimates of the mechanistic model parameters, especially when the number of unknown measurement mappings is low.

**Figure 6:**
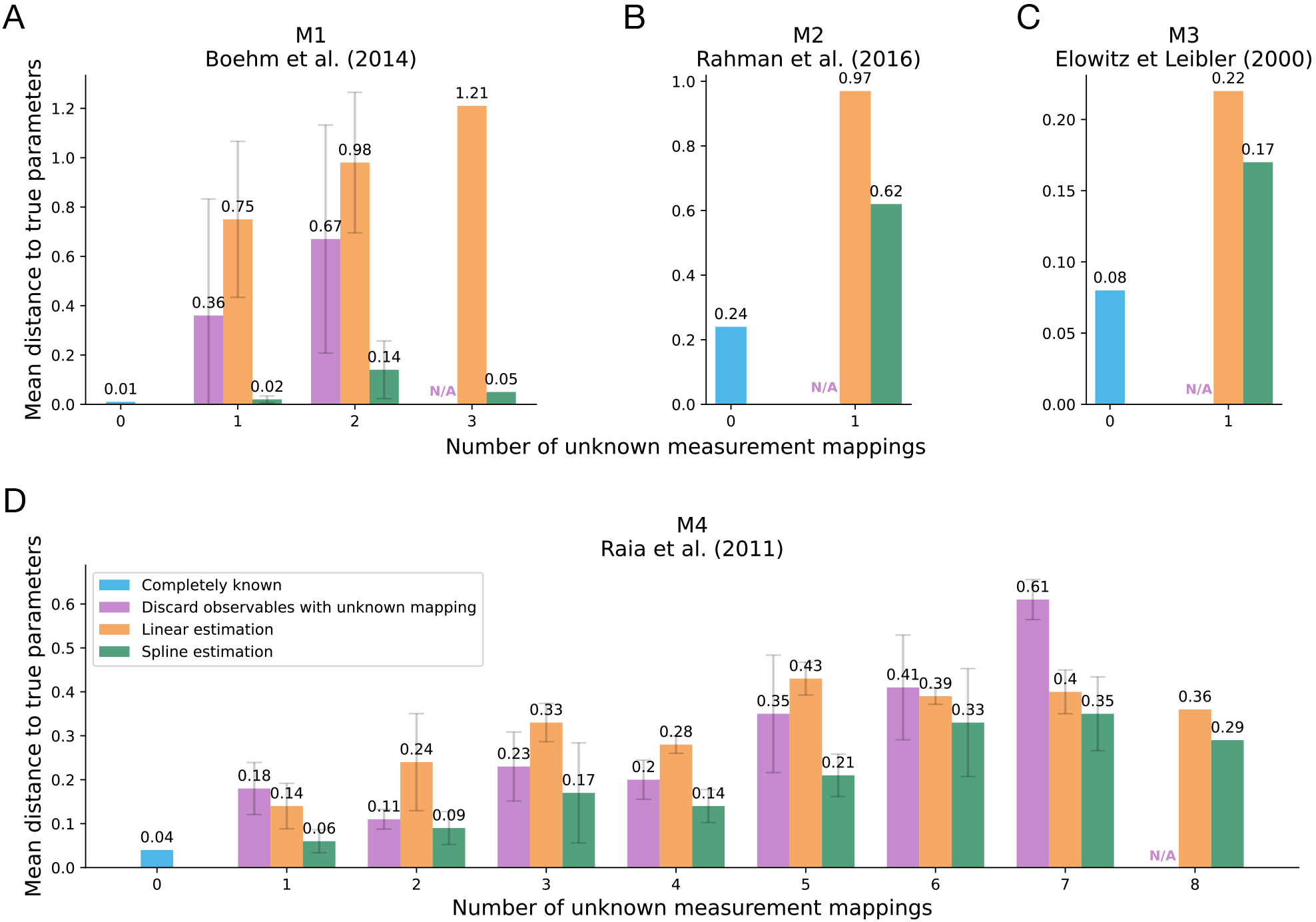
Evaluation of parameter inference across the number of unknown measurement mappings for linear and spline estimation. The parameter inference is measured by the L2 distance of the estimated to the true mechanistic model parameters. On the x-axis of each plot, we mark the model variant with a certain number of unknown measurement mappings, ranging from 0 to the number of model observables. The distance for each model variant is normalized by the number of mechanistic parameters and averaged across combinations of known-unknown observables. The best-case scenario for each model M1 - M4 is the case of completely known measurement mappings (**blue**). We compare the distance to the true parameters for the linear (**orange**) and spline (**green**) approach for each number of unknown measurement mappings. The approach of discarding the data of observables with unknown measurement mappings (**purple**) is depicted for reference. This approach is not feasible for the model variants with a maximum number of unknown measurement mappings, as that would involve the removal of all data, so we denote this with N/A.

## Discussion

Semi-quantitative measurements represent a large portion of the available data that can be used to estimate unknown mechanistic parameters of ODE models. Yet, there is a lack of a method to exploit this data, unless the mapping is linear. Here, we address this challenge by introducing a spline-based method for the estimation of unknown non-linear mappings. The approach can be applied to models with quantitative data for which it is unclear whether the data are truly linear (Fig. 7). Depending on the estimated optimal splines, the data can be deemed to be quantitative, relative, censored, or semi-quantitative, so that an appropriate method can be used. If the estimated optimal spline is non-linear, one can choose to estimate a parameterized function of a similar shape, or continue using the optimal splines as the measurement mapping. However, one has to be conservative with the number of observables chosen for spline estimation, as it can lead to a large expansion of the parameter space dimension. We evaluated the reliability of this process using an example, showing consistent qualitative measurement mapping shapes. An obvious extension of this approach is the inclusion of symbolic function identification from the estimated optimal splines. This would constitute an automatic parameterization of the unknown measurement mappings.

**Figure 7:**
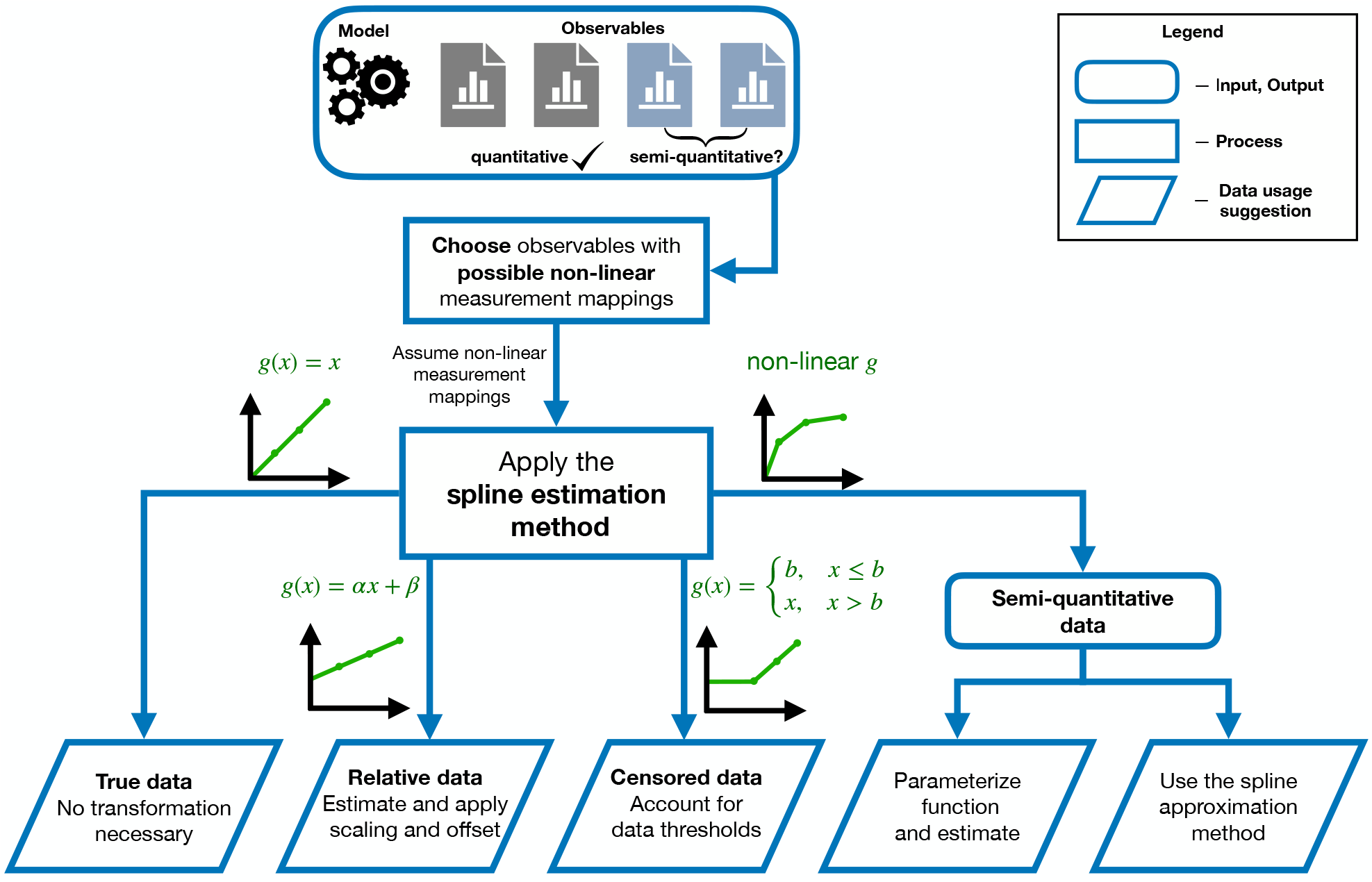
Diagram of application of the spline method to data with possible non-linear measurement mappings. Estimated optimal splines are depicted in green. Each arrow from the central square is a possible outcome of the non-linear mapping estimation.

To increase the method’s efficiency, we employed a hierarchical gradient-based optimization approach. We evaluated its performance and compared it with alternative approaches for four published models with differences in their complexity. This revealed a higher computational efficiency across all models. Further optimization acceleration could be achieved by including adjoint sensitivity analysis (ASA) [6, 14]. Even though our inner problem is not solved exactly, its gradient contribution is still zero. Thus, existing ASA software implementations from [22] can be used, since the gradient computation is the same as in hierarchical optimization with an exactly solved inner problem. Complementary to this, the derivation of second-order derivatives could further improve the method’s convergence and, with it, its computational efficiency.

The proposed method employs piece-wise linear splines to estimate general non-linear mappings. This was the simplest first-pass option, but, as they are not smooth, they had unavoidable approximation errors. Therefore, it is valuable to explore alternative smooth and flexible parameterized functions. Furthermore, they should retain the convexity of the inner optimization problems and the possibility of analytical gradient calculation. Interesting candidates are the scaled cumulative distribution functions (CDFs) of the beta distribution. They are monotone by definition, parameterized by only three parameters, and with promising flexibility to be able to model most types of measurement non-linear mappings.

In conclusion, we developed and implemented an easy-to-use framework to uncover unknown non-linear measurement mappings and to integrate semi-quantitative data into the parameter estimation of ODE models. The approach has a user-friendly implementation in the open-source Python Parameter Estimation TOolbox (pyPESTO). As it is agnostic to the structure of the underlying dynamical model, the method can be applied to models from different research fields, such as physics and engineering.

## Implementation and data availability

The proposed method is implemented in the open-source Python Parameter Estimation TOolbox (pyPESTO) [26]. Models M1-M4 were taken from the PEtab benchmark collection [21] based on [10]. For ODE integration, we used the AMICI Python toolbox [8]. Gradient-based optimization was performed using the fides optimizer [7] and gradient-free optimization was performed with the SciPy Powell algorithm [12]. Both optimizers were used through the pyPESTO interface with the default optimizer settings. All the code and models used in this study are available from the Zenodo database at https://doi.org/10.5281/zenodo.10568951.

## Author contributions statement

J.H. conceived the project. D.D. implemented the proposed approach and conducted all studies. S.G. and J.H. provided critical feedback on the implementation development. D.D. and J.H. wrote the manuscript. All authors discussed the results and commented on the manuscript.

## A Supplementary information

### A.1 Derivation of analytical gradient formulas

Here, we derive the formulas for the calculation of the analytical gradient of the objective function with respect to the mechanistic parameters in hierarchical optimization.

#### A.1.1 Model and spline definition

We consider models based on a system of ODEs

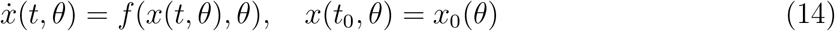

in which 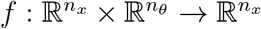 is a vector field describing the temporal evolution of the state variables 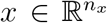, and 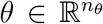 are the unknown mechanistic parameters. The measured properties of the model are its observables 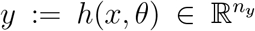. The dimensionalities of the state, parameter, and observable vector are denoted by *n*_*x*_, *n*_*θ*_ and *n*_*y*_, respectively. To simplify the following computations, we assume that our model has only one observable, that is, *n*_*y*_ = 1. Later in Subsection A.1.5, we show that this can be done without loss of generality. Additionally, let this observable be a non-linear semi-quantitative observable. In other words, there exists a true and unknown non-linear monotone mapping *g* : ℝ → ℝ such that, assuming an additive normally distributed noise model, the measured data 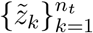 are given by:

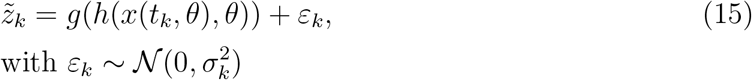

for time points 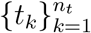, in which *n*_*t*_ is the number of observable time-points and 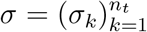 are the noise parameters. Noise parameters are also unknown in some cases and have to be estimated. In the case of such an observable, we refer to *g*(*h*(*x*(*t*_*k*_, *θ*), *θ*)) as the observable values, and to *y*_*k*_(*θ*) = *h*(*x*(*t*_*k*_, *θ*) as the (biochemical, epidemiological, …) quantities of interest. We approximated this non-linear mapping using a piecewise linear spline 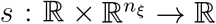, where *n*_*ξ*_ is the number of spline knots. The spline is given by

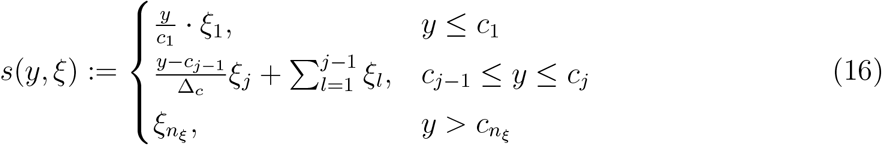

in which *ξ* are the differences between the heights of neighboring spline knots (Figure 8). In other words, the height of the *i* -th spline knot is equal to 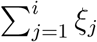. With 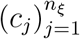, we denote the uniformly distributed bases of the spline knots, that is, *c*_*j*_ = *c*_1_ + (*j* − 1) · Δ_*c*_. In application, the true range of the biochemical or epidemiological quantities of interest is generally not known a priori. Thus, it is impossible to correctly fix the spline interval 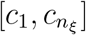. However, since we will use iterative optimization methods, at each optimization iteration, we will have access to the simulated quantities of interest 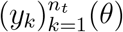. Therefore, to avoid this problem, we always scale the spline interval to the current model output interval 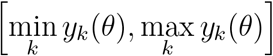. This makes the spline parameter bases dependent on the quantities of interest.

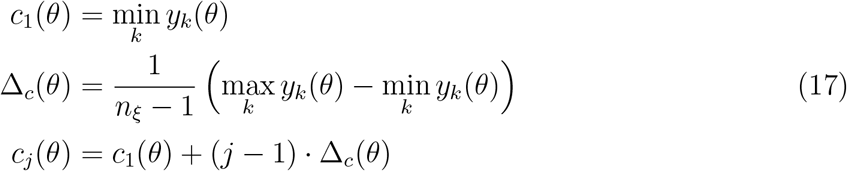

However, this does not cause any issues other than slightly increasing the complexity of the analytical gradient computation.

**Figure 8:**
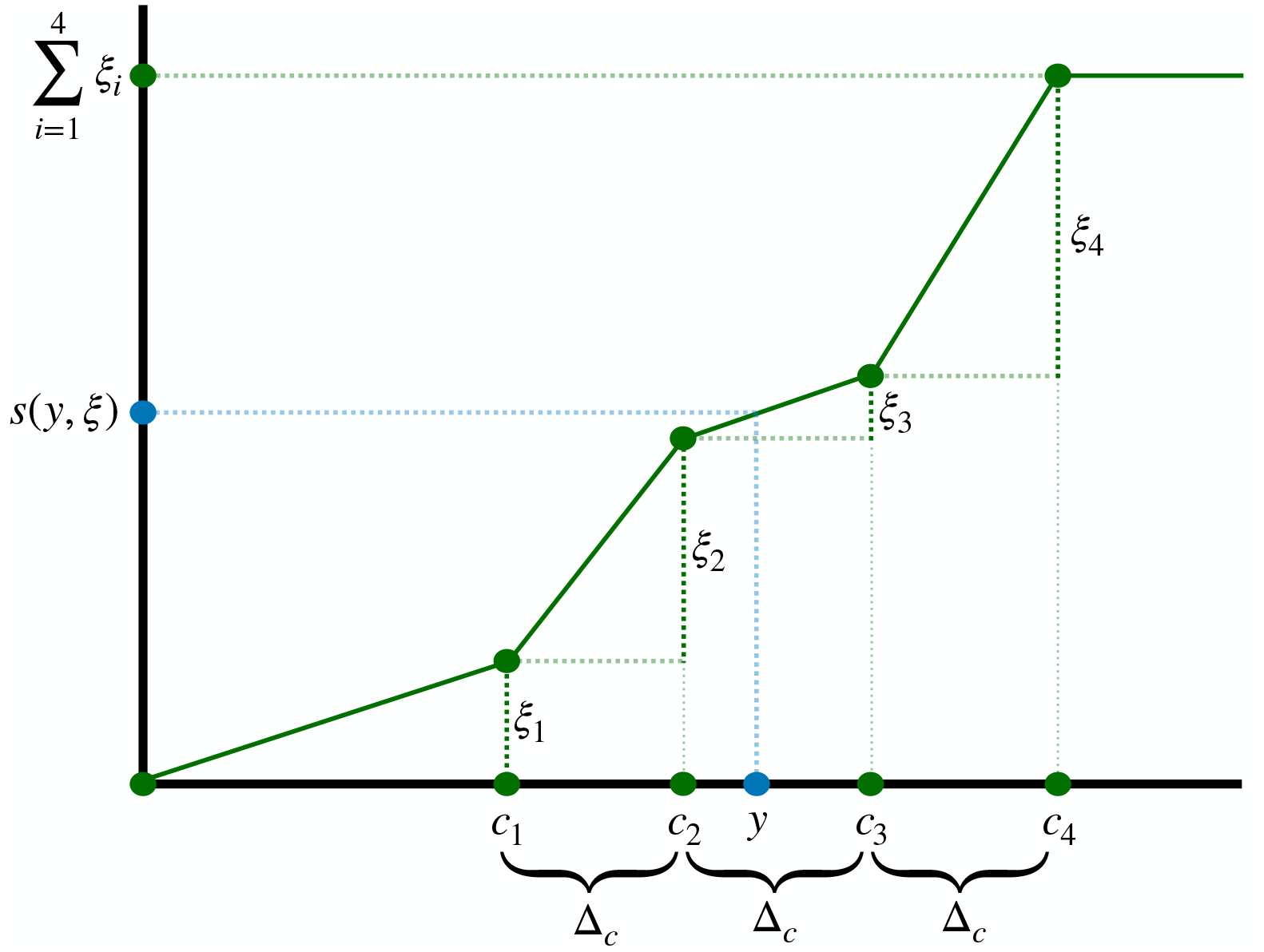
Illustration of the spline definition notation.

**Figure 9:**
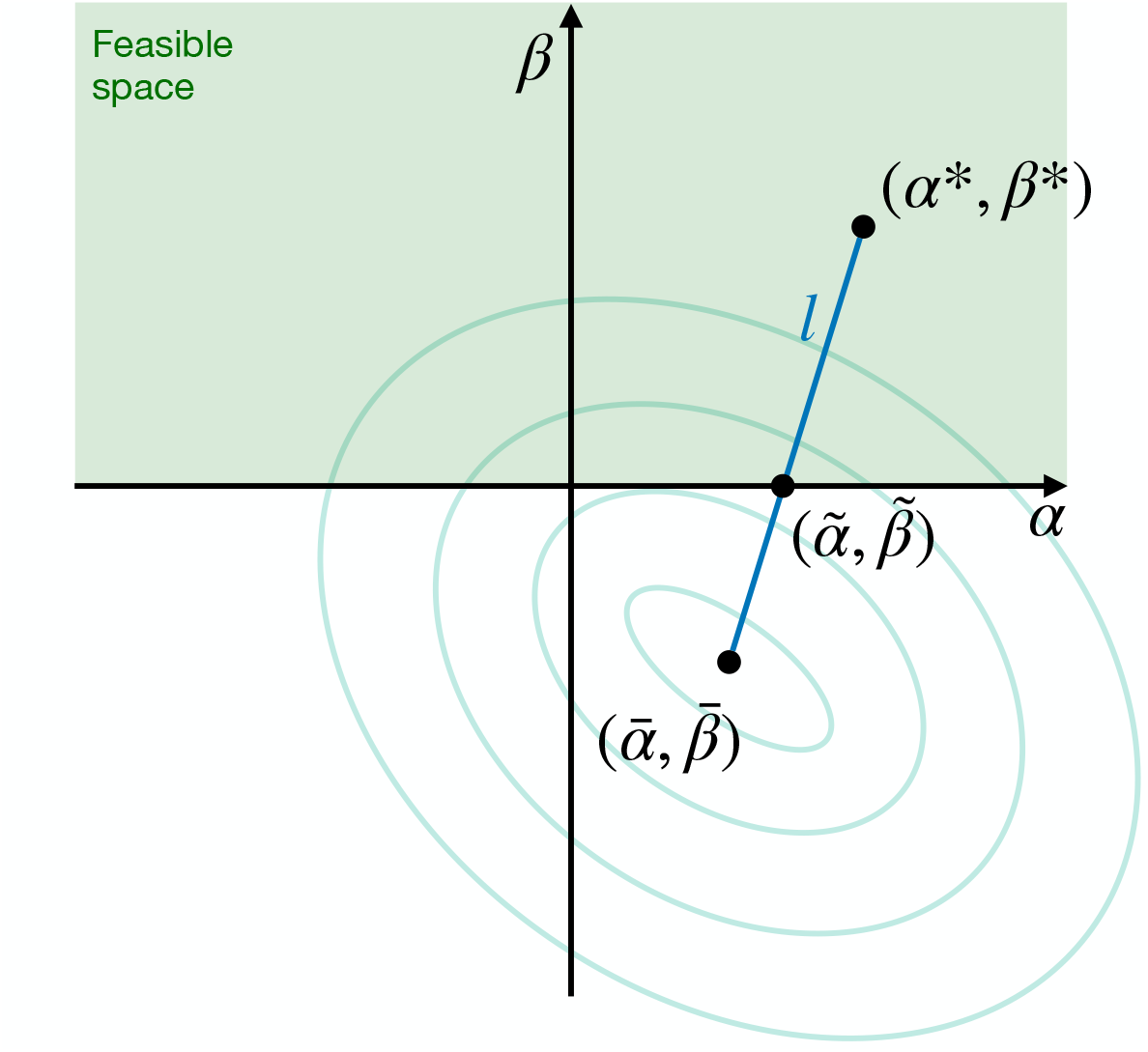
Proposition figure.

#### A.1.2 Objective function and the hierarchical optimization problem

The spline links the measured quantities to the quantities of interest of the non-linear measured observable

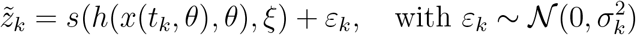

and thus yields the negative log-likelihood function

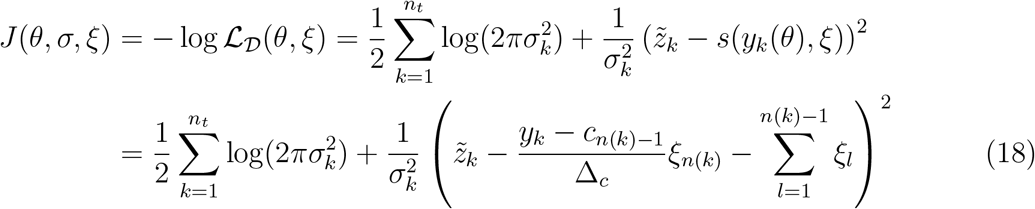

for a dataset 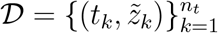 and simulated observables *y*_*k*_ = *h*(*x*(*t*_*k*_, *θ*), *θ*), for *k* = 1, …, *n*_*t*_. Here, 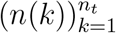 denote subinterval indices for simulated quantities of interest, i.e. *y*_*k*_ ∈ [*c*_*n*(*k*)−1_, *c*_*n*(*k*)_] for *k* = 1, …, *n*_*t*_. We note that currently the objective function depends on the mechanistic parameters *θ* only through simulations *y*_*k*_(*θ*). The objective function is optimized hierarchically in three nested levels:

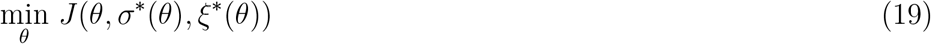

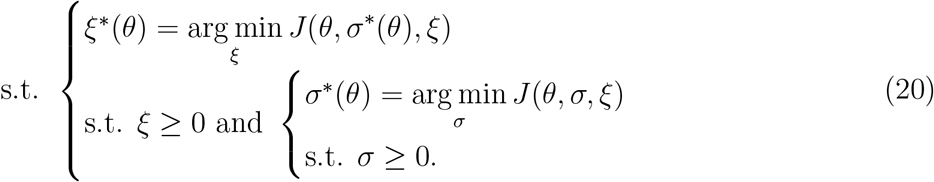

in which the mechanistic parameters are optimized in the outer optimization problem (19), whereas the spline parameters and noise parameters are optimized in two nested inner optimization problems (20). The true measurement mapping is assumed to be monotone, so the spline parameters are constrained to be positive. This is sufficient even in the case of a decreasing true measurement mapping, as inverting the sign of the quantities of interest will turn it into an increasing one. The main benefit of the hierarchical optimization framework is the simplicity of the inner problems. The inner optimization problem for noise parameters *σ* can be solved analytically, yielding

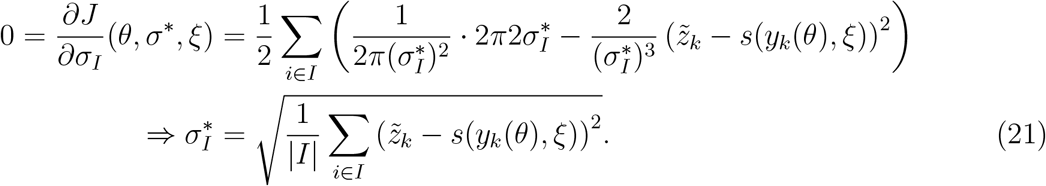

in which by *I* ⊂ {1, …, *n*_*t*_} we denote the subset of indices that share the noise parameter *σ*_*I*_. The *σ* non-negativity constraint is satisfied as this analytical result already provides a non-negative value. The inner problem for the spline parameters cannot be solved analytically, but we prove that it is convex in the following theorem. Therefore, its numerical optimization is computationally efficient.

We claim that the defined objective function is convex. We will prove this using the following theorem on the convexity of any least-squares optimization problem with non-negativity constraints.

##### Theorem 1

*Let an objective function* 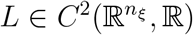 *be of the form*

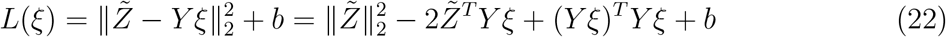

*Where* 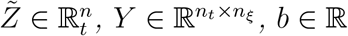, *and* ∥·∥_2_ *is the L2 norm. Then the optimization problem*

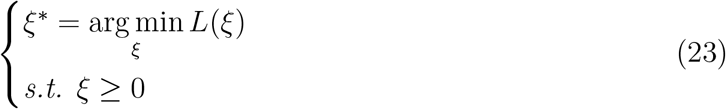

*is convex*.

*Proof*. We show that the Hessian of this function is positive semi-definite. The gradient of the objective function (22) is

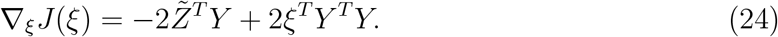

Hence, the Hessian matrix of the objective function at the point *ξ* is given by the Jacobian of the gradient

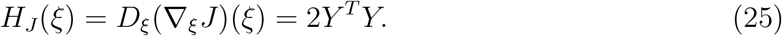

This is a positive semi-definite matrix since, for any vector 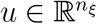, the following holds:

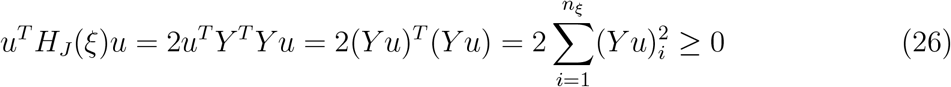

where (*Y u*)_*i*_ is the *i*-th element of the vector *Y u*. In combination with affine inequality constraints, this implies that the optimization problem is convex.

The negative log-likelihood function defined in (18) can be easily reformulated into the form (22), where 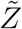, *Y*, and *b* can depend on *θ* and *σ* but not on *ξ*. Thus, the inner optimization problem for the spline parameters is convex.

#### A.1.3 Analytical gradient

Due to the dependency of the optimal spline parameters *ξ*\* and optimal noise parameters *σ*\* on the mechanistic parameters *θ*, the derivative of the objective function with respect to a mechanistic parameter *θ*_*i*_ contains two additional terms

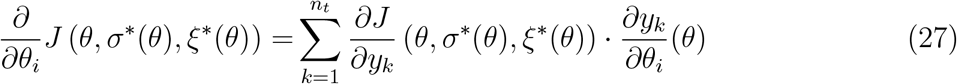

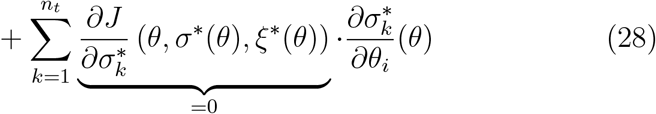

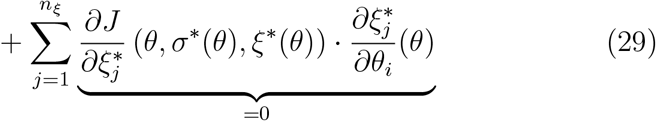

where we refer to 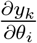 as the observable sensitivity at time point *t*_*k*_ with respect to the parameter *θ*_*i*_. The first gradient term can be calculated using forward sensitivity analysis (FSA) or adjoint sensitivity analysis (ASA). Furthermore, the inner problem with respect to the noise parameters is solved exactly. Therefore, the second term (28) is always 0. This leaves the third term to be obtained. In the following theorem, we prove that it is always equal to 0 as well. The theorem is equivalent to the envelope theorem, which is used mainly in economic theory [3].

##### Theorem 2

*For any* 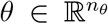, *let the pair* 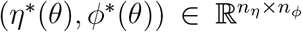 *be a solution of the following optimization problem*

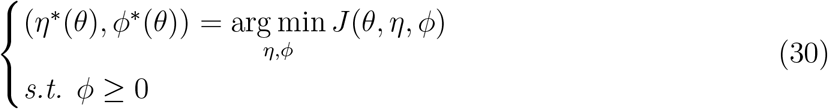

*where* 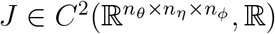 *is a convex objective function and ϕ is constrained to be positive. Then for all* 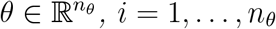 *and j* = 1, …, *n*_*ϕ*_, *k* = 1, …, *n*_*η*_

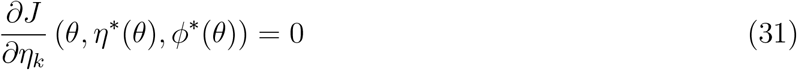

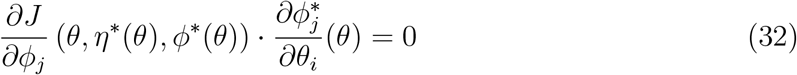

*Proof*. The pair (*η*\*(*θ*), *ϕ*\*(*θ*)) is a solution to a convex optimization problem with affine inequality constraints. Therefore, Slater’s condition holds [28], and there exist optimal Lagrange multipliers 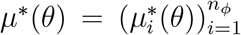 such that the triplet (*η*\*(*θ*), *ϕ*\*(*θ*), *µ*\*(*θ*)) satisfies the Karush-Kuhn-Tucker (KKT) conditions

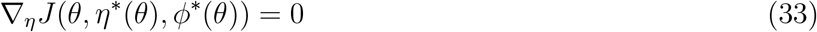

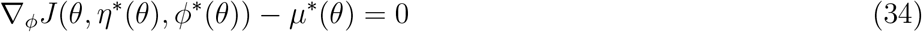

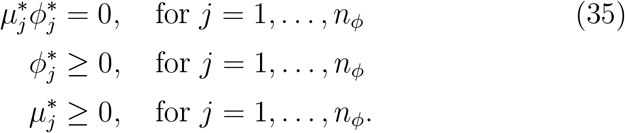

The first KKT condition (33) directly proves the first statement of the Theorem (31). Furthermore, due to the simple positivity constraints, the second KKT condition (34) states that the Lagrange multipliers are equal to the objective function gradient with respect to *ϕ*

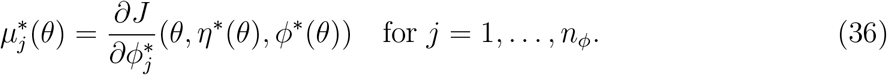

To prove the first statement of the theorem (31) for an index *j* ∈ {1, …, *n*_*ϕ*_}, we consider two cases: 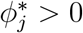 and 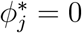.

Case 1: 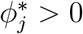. From the third KKT condition (35), we then know that 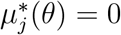. Therefore, the statement follows from (36).

Case 2: 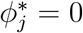. For an index *i* ∈ {1, …, *n*_*θ*_}, we differentiate the third KKT condition (35) with respect to *θ*_*i*_ to obtain

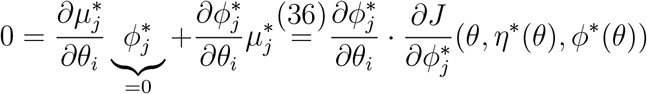

□

Since the negative log-likelihood function defined in (18) is convex it satisfies the theorem’s conditions. In addition, in this case, *η* = *σ* and *ϕ* = *ξ*. Therefore, the derivative of the objective function with respect to a mechanistic parameter *θ*_*i*_ can be analytically computed and is equal to

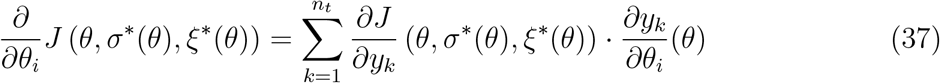

#### A.1.4 Spline regularization

In our implementation, the spline (16) is additionally regularized to reduce overfitting. The regularization penalizes the non-linearity of the spline with the aim of minimizing false-positive non-linear measurement mappings. To achieve this, we add the following regularization term to the objective function:

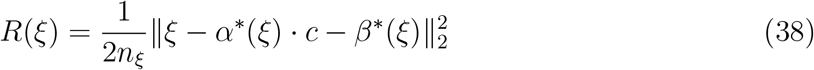

where

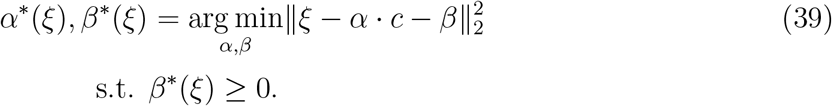

In other words, *α*\*(*ξ*) and *β*\*(*ξ*) are the optimal scaling and offset of a linear regression of the spline knots 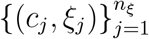 and the regularization is a penalization of the distance between the spline knots and the optimal linear line. Having a negative offset is not plausible in most applications. Thus, we constrain the offset to be non-negative.

We will show this optimization problem can be solved analytically. To do so, we state and prove the following simple proposition.

##### Proposition

*Let* 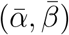 *be the unique solution of an unconstrained optimization problem*

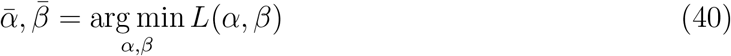

*where L* : ℝ ^2^ → ℝ *is a strictly convex objective function. Then the unique solution* (*α*\*, *β*\*) *of the optimization problem with a positivity constraint*

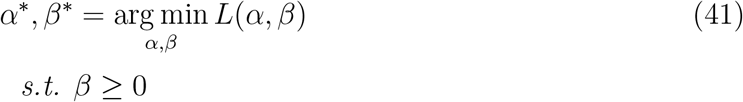

*satisfies the following:*

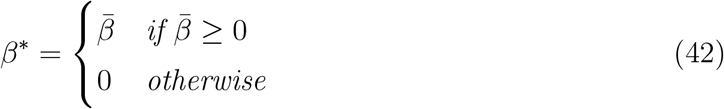

In other words, we can solve for the positivity-constrained variable by simply solving the unconstrained problem and inspecting whether it is positive. If it is, then the solution is given by it. Otherwise, the optimal solution is on the constraint, i.e. 0.

*Proof*. The first statement of the proposition is trivial. If the solution of the unconstrained problem lies in the feasible region of the constrained problem, then the solutions 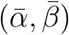 and (*α*\*, *β*\*) are equal due to the same strictly convex objective function of the optimization. To prove the second statement, let us assume the opposite: 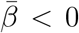, but *β*\* *>* 0. Then we connect the two solutions by a line in the two-dimensional parameter space *l* : [0, 1] → ℝ ^2^ from (*α*\*, *β*\*) to 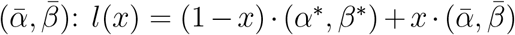. Since these points are on opposite sides of the constraint, the line will cross the constraint at some point 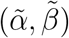. This point also satisfies 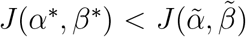 because it lies in the feasible parameter space and is not equal to the optimal point (*α*\*, *β*\*) due to being on the constraint 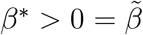. However, from the solution of the unconstrained problem, we know that 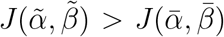. In conclusion, we have found a line *l* with three points on it that satisfy 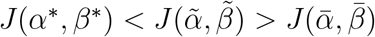. This is in contradiction to the strict convexity of the function *J* and therefore the second statement holds as well.

Now we will apply the proposition to our problem. The optimization problem (39) satisfies the conditions of Theorem (1), so it is convex. Furthermore, it is strictly convex due to the full rank of the corresponding Hessian matrix *Y* ^*T*^ *Y*, which makes the inequality (26) strict. Thus, it satisfies the conditions of the proposition. To obtain the solution of the unconstrained version of the problem, we equate the gradient of the objective function to 0.

This gives a linear system whose solution is

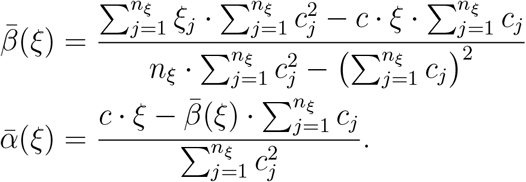

According to the proposition, to obtain the solution to the unconstrained problem we inspect the sign of 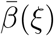. If it is positive, the solution is equal to 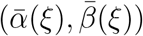. Otherwise, the optimal *β*\*(*ξ*) is equal to 0. To obtain the optimal scaling *α*\*(*ξ*) for this case we turn to the KKT conditions of the constrained optimization problem:

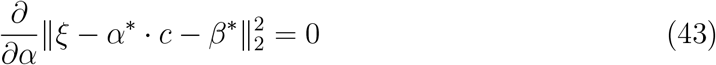

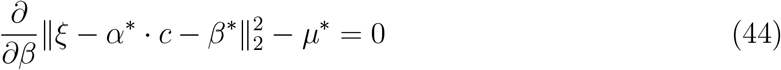

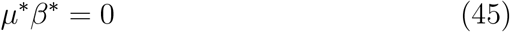

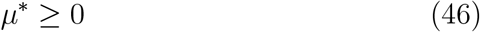

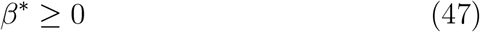

where *µ*\* ∈ ℝ is the optimal Lagrange multiplier. The first KKT condition (43) is a linear equation in *α*\*(*ξ*) whose solution for *β*\*(*ξ*) = 0 is

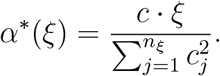

Pulling the two cases together, we can write the final analytical solution to the optimization problem as

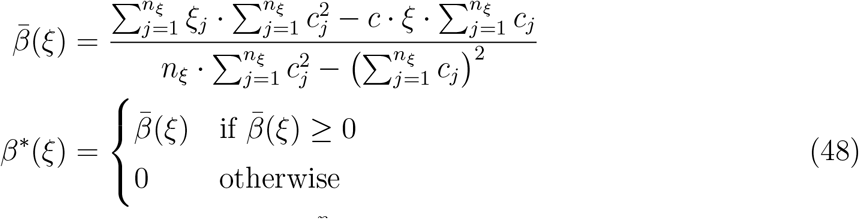

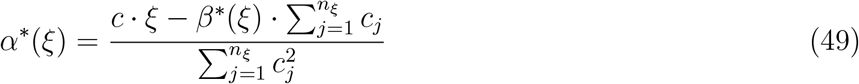

This regularization is appended to the objective function of the model optimization (18). Thus, the complete objective function is given as:

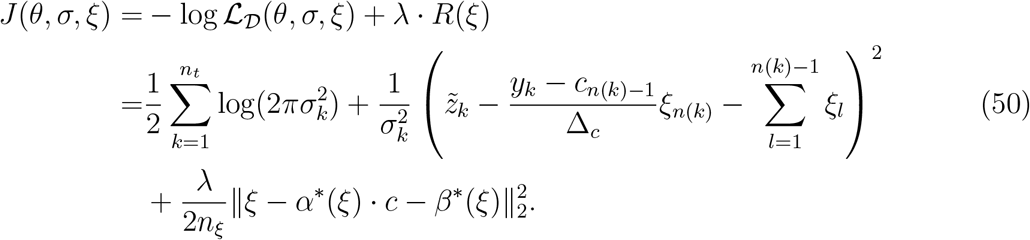

where *λ* ∈ ℝ^+^ is the regularization strength parameter. We note that this addition does not change the result of the analytical gradient (Theorem 2), as the objective function remains convex.

#### A.1.5 Models with multiple observables

Here, we show that the results obtained in previous subsections hold for a model with an arbitrary number of semi-quantitative observables *n*_*y*_. The semi-quantitative data is linked to the observables via

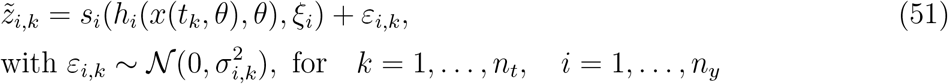

where each semi-quantitative observable has its own spline function 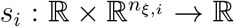. We assume that each one of the splines depends on its own vector of spline parameters *ξ*_*i*_ and that these parameters are not shared among observables. Similarly, we assume that the observable noise parameters *σ*_*i*_ are not shared between observables. Thus, the overall objective function *J* consists of *n*_*y*_ observable-specific objective functions 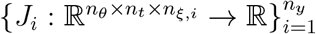

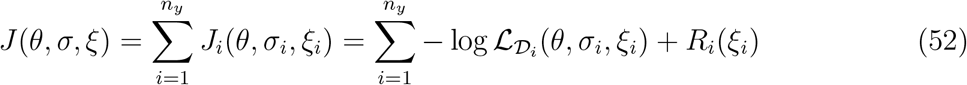

where 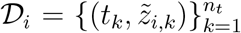 is the data set of the *i*-th observable. Therefore, the hierarchical optimization problem contains *n*_*y*_ separate inner sub-problems that can be solved as before.

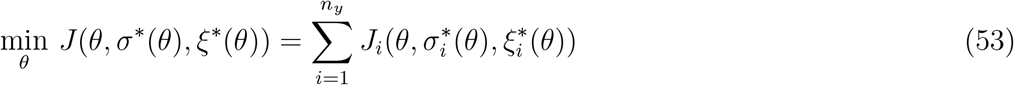

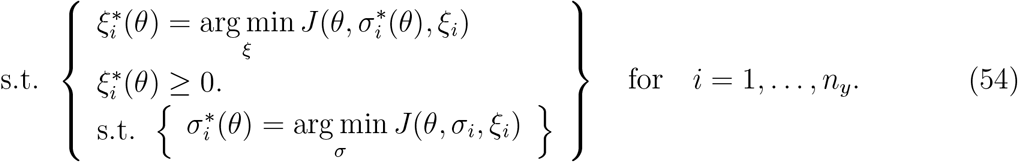

To efficiently solve the outer optimization problem, we need to calculate the gradient of the objective function *J* with respect to the mechanistic parameters *θ*. Since the objective function is additive in observables, this is given by:

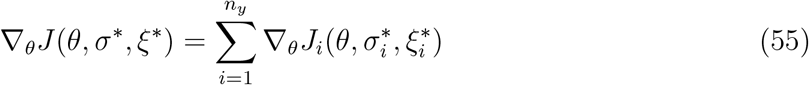

where 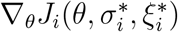 can be obtained as before.

### A.2 Model details

For the evaluation of the proposed method, we employed five models in total: one toy model T1 and four published models M1-M4. The models contain a varying number of states, mechanistic parameters, observables, and data points (Table 2). The published models are taken from a collection of parameter estimation problems in PEtab format, which is based on the benchmark collection by [10].

**Table 2:** Application models. Consistent with notation from Section A.1, *n*_*x*_, *n*_*θ*_, and *n*_*y*_ denote the numbers of state variables, mechanistic parameters, and observables, respectively. With |*𝒟*| we denote the cardinality of the data set.

| Model | $n_x$ | $n_\theta$ | $n_y$ | $\vert \mathcal{D}\vert $ | Description | Reference |
| --- | --- | --- | --- | --- | --- | --- |
| T1 | 2 | 4 | 1 | 12 | FRET probe activation | [1] |
| M1 | 8 | 6 | 3 | 48 | STAT5 dimerization | [2] |
| M2 | 7 | 9 | 1 | 23 | Infectious disease dynamics | [19] |
| M3 | 8 | 18 | 1 | 58 | Transcriptional regulation | [4] |
| M4 | 14 | 18 | 8 | 205 | IL13-induced signaling | [20] |

These models were originally calibrated on quantitative measurements. Thus, to benchmark the proposed method, we modified the observation model to include non-linear measurement mappings. However, as the original model measurements were already noise-corrupt, it is not possible to retain the same noise model by simply applying the non-linear mappings to those measurements. Therefore, we generated synthetic noise-free data and transformed it with the non-linear mappings. Only then could we add noise of the same original magnitude. This resulted in the same noise model as in the original model versions. The mechanistic parameters used to generate the synthetic data were estimated using the original quantitative measurements. These nominal mechanistic parameters are available in the benchmark collection mentioned above.

#### A.2.1 Model T1: FRET probe activation

The toy model is a simple model of FRET probe activation introduced by [1]. The model assumes Michaelis-Menten (MM) mechanisms for both probe activation and deactivation. All mechanistic, noise, and observable parameters are presented in Table 3. The model consists of one differential equation and one conservation law:

**Table 3:**
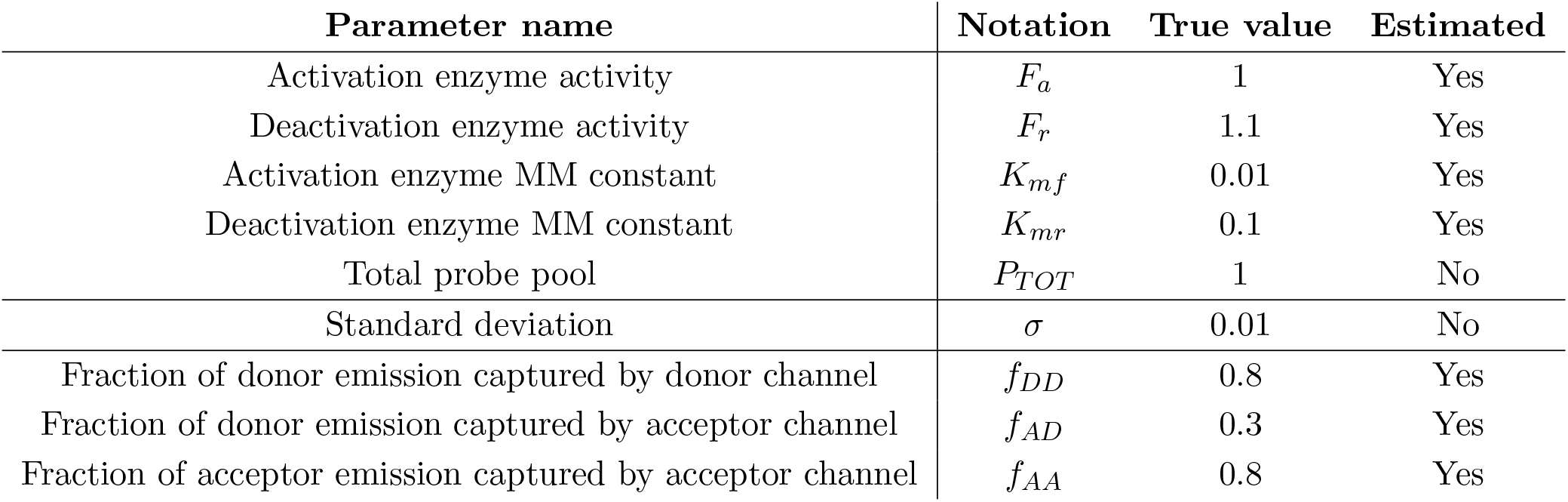
Toy model T1 parameters. The horizontal lines separate mechanistic, noise, and observable parameters, respectively.

| Parameter name | Notation | True value | Estimated |
| --- | --- | --- | --- |
| Activation enzyme activity | $F_a$ | 1 | Yes |
| Deactivation enzyme activity | $F_r$ | 1.1 | Yes |
| Activation enzyme MM constant | $K_{mf}$ | 0.01 | Yes |
| Deactivation enzyme MM constant | $K_{mr}$ | 0.1 | Yes |
| Total probe pool | $P_{TOT}$ | 1 | No |
| Standard deviation | $\sigma$ | 0.01 | No |
| Fraction of donor emission captured by donor channel | $f_{DD}$ | 0.8 | Yes |
| Fraction of donor emission captured by acceptor channel | $f_{AD}$ | 0.3 | Yes |
| Fraction of acceptor emission captured by acceptor channel | $f_{AA}$ | 0.8 | Yes |

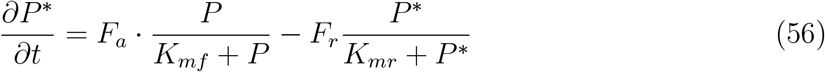

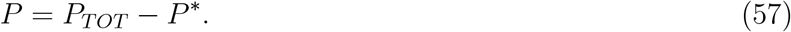

Furthermore, the model’s only observable is the ratiometric imaging intensity ratio

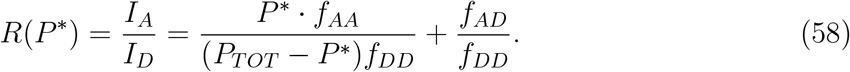

Consistent with the notation established in Section A.1, the observation function of this observable is the simple identity function *h*(*P*\*) = *P*\*, whereas the non-linear measurement mapping *g* is given by *R*.

To obtain synthetic data, we simulated the model with the true values of mechanistic parameters from Table 3, applied the measurement mapping function (58), and added additive normal noise with standard deviation given in the same Table.

Three model variants with varying observable models were estimated. The first is the parameterized model. Its observation model was a parameterization of the true measurement mapping (58). To avoid non-identifiability of observable parameters, instead of directly estimating *f*_*AA*_, *f*_*AD*_ and *f*_*DD*_, we rather estimated a parameterization with only two observable parameters *α* and *β*

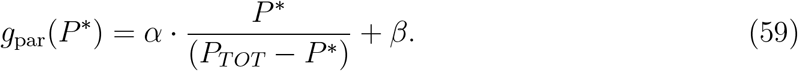

The second model estimated the measurement mapping using a linear function. Lastly, the third model used the proposed spline estimation method.

#### A.2.2 Model M1: STAT5 dimerization

The STAT5 dimerization model was introduced by [2]. It possesses 3 observables – pSTAT5A rel, pSTAT5B rel, and rSTAT5A_rel. We will refer to the observables as first, second, and third in that order. The chosen synthetic non-linear measurement mappings are shown in Figure 10: saturation, sigmoid, and censoring. In the same order, their expressions are

**Figure 10:**
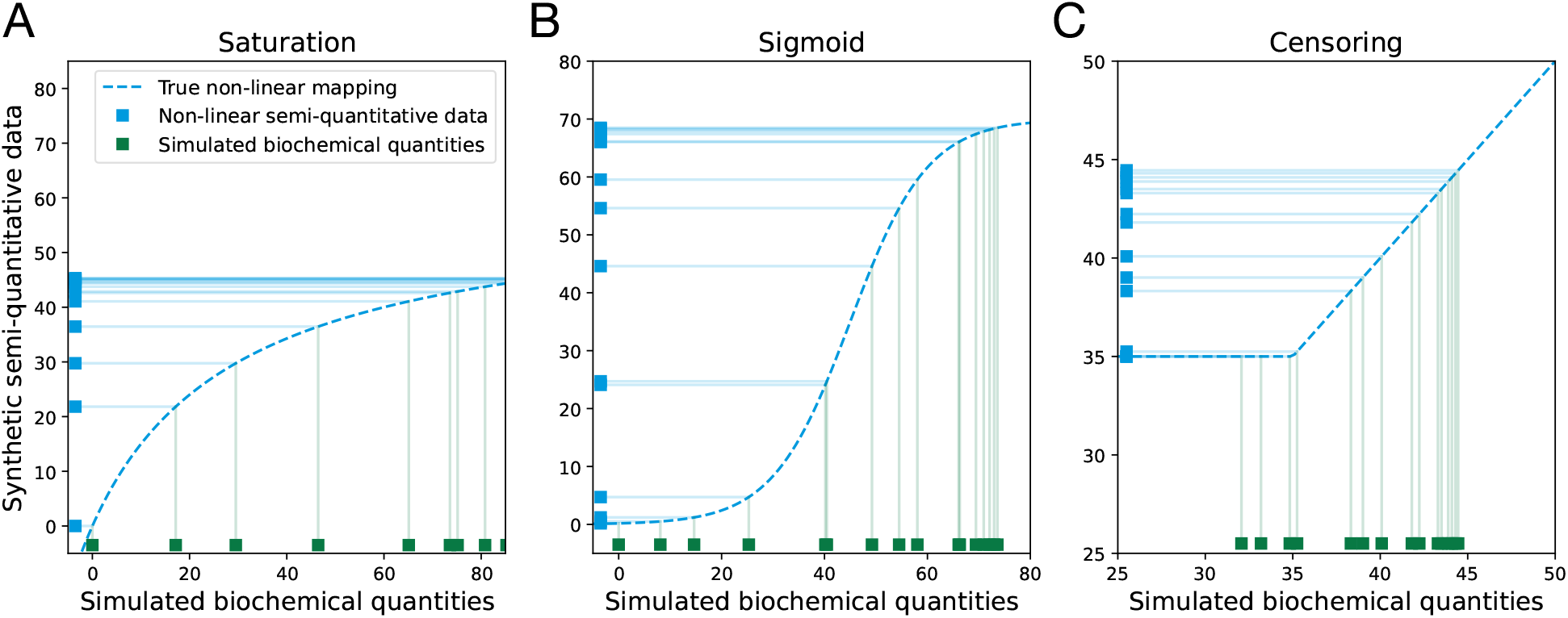
Synthetic non-linear semi-quantitative data of model M1. The simulated quantities of interest are denoted on the x-axis in green, whereas the semi-quantitative measurements are denoted on the y-axis in blue. The chosen non-linear measurement mappings are denoted in dashed blue for each observable (A-C).

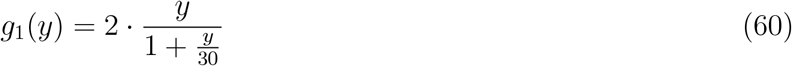

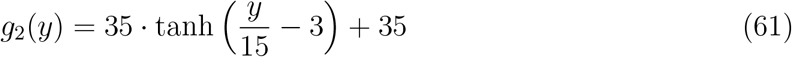

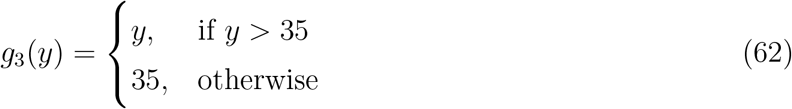

where the specific numerical values were chosen such that the mappings are significantly non-linear in the range of the respective simulated biochemical quantities.

#### A.2.3 Model M2: Infections disease dynamics

The model of infectious disease dynamics was introduced by [19]. It describes the impacts of early treatment programs on HIV epidemics and overall community-level immunity. It possesses only one observable, HIV prevalence. The chosen non-linear measurement mapping is a hyperbolic growth function

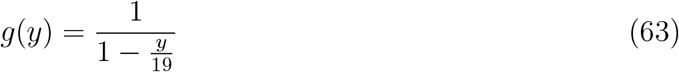

where the specific parameterization was chosen such that the mapping is significantly non-linear in the range of the simulated quantity of interest.

#### A.2.4 Model M3: Transcriptional regulation

The model of transcriptional regulation was introduced by [4]. It models an oscillating network consisting of three transcriptional repressor systems, termed the repressilator, in Escherichia coli. It possesses only one observable – protein fluorescence readout. The chosen non-linear measurement mapping is a saturation function

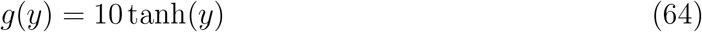

where the specific parameterization was chosen such that the mapping is significantly non-linear in the range of the simulated quantity of interest.

#### A.2.5 Model M4: IL13-induced signaling

The model of IL13-induced JAK2/STAT5 pathway signaling in lymphoma cell lines was introduced by [20]. It possesses 8 observables. We chose non-linear measurement mappings of the same shape across observables, scaled to the range of the simulated quantities of interest. The functions are

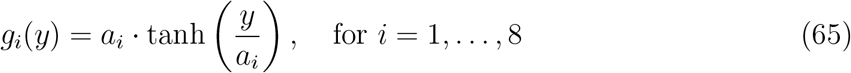

Where 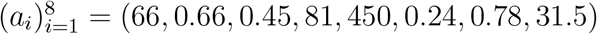 are the scaling factors chosen such that the mappings are significantly non-linear in the range of the respective simulated quantities of interest.

### A.3 Implementation and optimization details

The proposed method is implemented in the open-source Python Parameter Estimation TOolbox (pyPESTO, [26]). The implementation allows for the specification of an arbitrary number of spline knots and spline regularization strength. Within the GitHub repository of pyPESTO, there is a Jupyter notebook available that showcases the usage of the method.

Parameter estimation of all published models was performed using multi-start local optimization with 1000 starts per model. Each estimation was run on 10 cores of the AMD EPYC 7F72 3.20 GHz processor with 1TB of RAM. Gradient-based optimization was performed using the fides optimizer [7] and gradient-free optimization was performed with the SciPy Powell algorithm [12]. Both optimizers were accessed through the pyPESTO interface with the default optimizer settings. For optimization of inner problems, we used SciPy’s L-BFGS-B algorithm with default settings. For ODE integration, we used the AMICI Python toolbox [8].

### A.4 Parameter inference study

In the last subsection of the Results section in the manuscript, we presented a study focused on parameter inference. We compared the linear estimation and spline estimation approaches in the four application examples M1 - M4 with their respective synthetic non-linear measurement mappings. Furthermore, we evaluated the impact of an increasing number of unknown measurement mappings.

For illustration, we consider the model M1. It contains three observables, each with its own non-linear measurement mapping (Fig. 10). However, since these mappings are synthetic and thus known, we were able to include the mappings 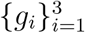 in the observable function 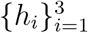, transforming the semi-quantitative observables into standard quantitative ones. This is how we obtained model variants of the same initial model M1 with a variable number of unknown measurement mappings. Furthermore, to remove any observable bias, we considered all combinations of unknown and known observables (Table 4).

**Table 4:**
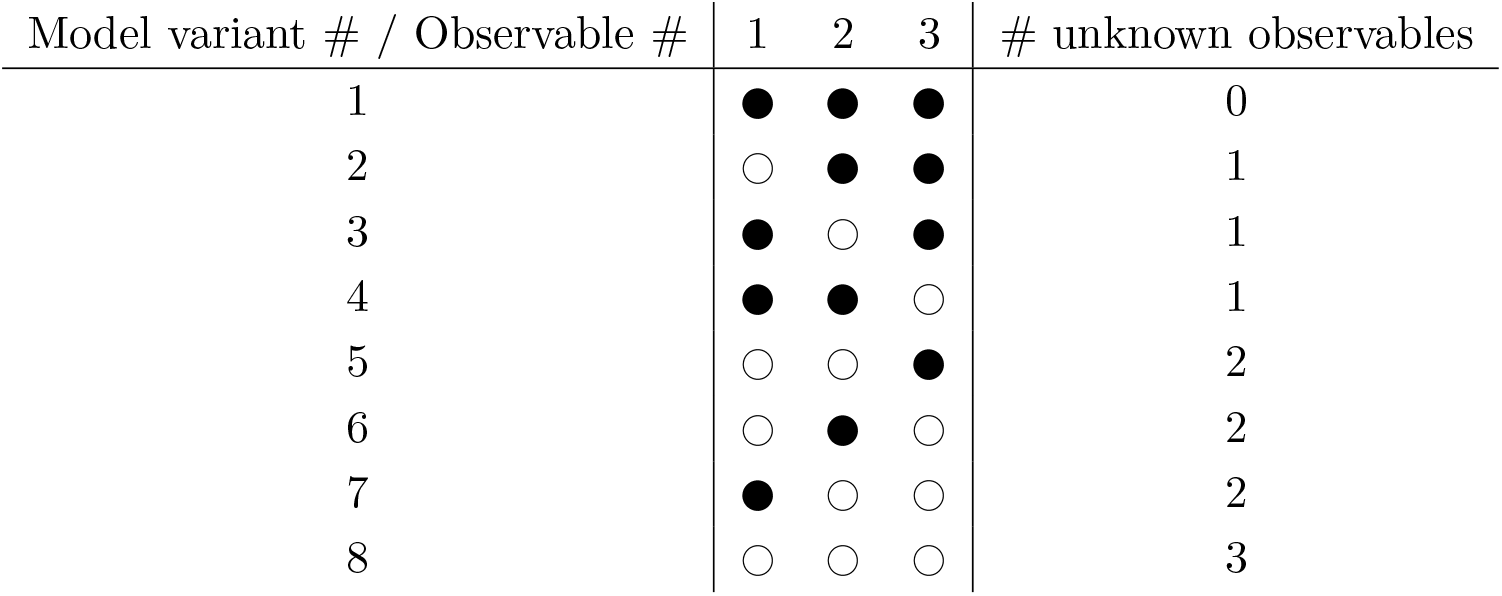
Observable combinations of model M1. ● and ○ denote whether the measurement mapping of the observable is known or unknown, respectively.

For models M2 and M3 this is simple, as they contain only one observable so we considered two variants: known observable mapping and unknown observable mapping. Model M4 contains 8 observables, so it would have been computationally too expensive to include all 2^8^ = 256 model variants with 3 estimation approaches. Therefore, to reduce the observable bias while keeping the number of model variants reasonably low, we considered the model M4 variants shown in Table 5.

**Table 5:** Observable combinations of model M4. ● and ○ denote whether the measurement mapping of the observable is known or unknown, respectively.

| Model variant # / Observable # | 1 | 2 | 3 | 4 | 5 | 6 | 7 | 8 | # unknown observables |
| --- | --- | --- | --- | --- | --- | --- | --- | --- | --- |
| 1 | ● | ● | ● | ● | ● | ● | ● | ● | 0 |
| 2 | ○ | ● | ● | ● | ● | ● | ● | ● | 1 |
| 3 | ○ | ○ | ● | ● | ● | ● | ● | ● | 2 |
| 4 | ○ | ○ | ○ | ● | ● | ● | ● | ● | 3 |
| 5 | ○ | ○ | ○ | ○ | ● | ● | ● | ● | 4 |
| 6 | ○ | ○ | ○ | ○ | ○ | ● | ● | ● | 5 |
| 7 | ○ | ○ | ○ | ○ | ○ | ○ | ● | ● | 6 |
| 8 | ○ | ○ | ○ | ○ | ○ | ○ | ○ | ● | 7 |
| 9 | ● | ● | ● | ● | ● | ● | ● | ○ | 1 |
| 10 | ● | ● | ● | ● | ● | ● | ○ | ○ | 2 |
| 11 | ● | ● | ● | ● | ● | ○ | ○ | ○ | 3 |
| 12 | ● | ● | ● | ● | ○ | ○ | ○ | ○ | 4 |
| 13 | ● | ● | ● | ○ | ○ | ○ | ○ | ○ | 5 |
| 14 | ● | ● | ○ | ○ | ○ | ○ | ○ | ○ | 6 |
| 15 | ● | ○ | ○ | ○ | ○ | ○ | ○ | ○ | 7 |
| 16 | ○ | ● | ● | ● | ● | ● | ● | ● | 1 |
| 17 | ○ | ● | ● | ● | ● | ● | ● | ○ | 2 |
| 18 | ○ | ○ | ● | ● | ● | ● | ● | ○ | 3 |
| 19 | ○ | ○ | ● | ● | ● | ● | ○ | ○ | 4 |
| 20 | ○ | ○ | ○ | ● | ● | ● | ○ | ○ | 5 |
| 21 | ○ | ○ | ○ | ● | ● | ○ | ○ | ○ | 6 |
| 22 | ○ | ○ | ○ | ○ | ● | ○ | ○ | ○ | 7 |
| 23 | ○ | ○ | ○ | ○ | ○ | ○ | ○ | ○ | 8 |

